# Biochemical and structural characterization of a Class A β-lactamase from *Nocardia cyriacigeorgica*

**DOI:** 10.1101/2023.10.02.560429

**Authors:** Jérôme Feuillard, Julie Couston, Yvonne Benito, Elisabeth Hodille, Oana Dumitrescu, Mickaël Blaise

**Affiliations:** IRIM, CNRS, Montpellier University, Montpellier, France; Institut des Agents Infectieux, Hospices Civils de Lyon, Hôpital de la Croix-Rousse, Centre de Biologie Nord, Lyon, France; Centre International de Recherche en Infectiologie (CIRI), INSERM U1111, CNRS UMR5308, ENS Lyon, Université Lyon 1, Lyon, France

**Keywords:** *Nocardia* spp., *Nocardia cyriacigeorgica*, class A ß-lactamase, ß-lactam resistance

## Abstract

*Nocardia* are gram-positive bacteria from the Actinobacteria phylum. Some *Nocardia* species can infect humans and are usually considered opportunist pathogens as they often infect immunocompromised patients. Albeit, their clinical incidence is low, many *Nocardia* species are nowadays considered emerging pathogens. Primary sites of infection of *Nocardia* are the skin or the lungs but dissemination to other body parts is very frequent. These disseminated infections are very difficult to treat and thus, tackled with multiple classes of antibiotics, on top of the traditional treatment targeting the folate pathway. ß-lactams are often included in the regimen but many *Nocardia* species present moderate or strong resistance to some of this drug class. Genomic, microbiological, and biochemical studies have reported the presence of class A ß-lactamases (ABL) in a handful of *Nocardia* species but no structural investigation of *Nocardia* ß-lactamases has been performed yet. In this study, we report the expression, purification, and preliminary biochemical characterization of the ABL from a *Nocardia cyriacigeorgica* (NCY-1) clinical strain. We describe, as well, the crystallization and the very high-resolution crystal structure of NCY-1. The protein sequence and structural analysis demonstrate that NCY-1 belongs to ß-lactamase of class A1 and attest to its very high conservation with ABL from other human pathogenic *Nocardia*. In addition, the presence of one molecule of citrate tightly bound in the catalytic site of the enzyme is described. This structure may provide a solid basis for future drug development to specifically target *Nocardia* spp. ß-lactamases.

**Synopsis:** The crystal structure at high resolution of a class A ß-lactamase from a clinical strain of *Nocardia cyriacigeorgica* is reported.

## 1 Introduction

Nocardiosis is an infectious disease caused by environmental bacteria from the *Nocardia* gender. *Nocardia* are Gram-positive bacteria from the Actinomycetota (or Actinobacteria) phylum. To date, about 130 species of *Nocardia* have been reported (https://www.bacterio.net/genus/nocardia) of which about 40 species are human pathogens and clinically relevant (Mehta and Shamoo, 2020). *Nocardia* are usually opportunistic bacteria that often trigger pulmonary infections. They also frequently disseminate to other organs, skins, joints, or the brain. Central nervous infections are very acute forms of nocardiosis as the mortality rate ranges from 40 to 80% (Cattaneo et al., 2013; McNeil and Brown, 1994; Wilson, 2012). The incidence of *Nocardia* infections is rather low but certainly underestimated as identification of *Nocardia* species has been impaired by technical aspects, as it is mainly based on gene sequencing. Powerful methods such as matrix-assisted laser desorption/ionization time-of-flight (MALDI-TOF) mass spectrometry (Hodille et al., 2023) will likely enable better identification and may help apprehend the real incidence of this emerging class of bacteria.

For long the reference treatment of nocardiosis, whatever the species, was based on a combination of two antibiotics, trimethoprim (TMP) and sulfamethoxazole (SMX), targeting the folic acid pathway (Margalit et al., 2021). Nonetheless, treating nocardiosis with this regimen remains very challenging and is often punctuated by treatment failure and/or relapse. This is mainly due to the high prevalence of drug resistance strains to TMP-SMX (Uhde et al., 2010). It is extremely difficult to treat the disseminated forms of *Nocardia* particularly those of the central nervous system. Treatment success is also impaired by the difficulty of choosing and adapting the right drug regimen since there is a large heterogeneity of drug susceptibility and drug resistance between the different human pathogenic *Nocardia* species (Hershko et al., 2023; Lebeaux et al., 2019).

To circumvent these issues, the current recommendation is therefore a combination of different antibiotic classes. On top of aminoglycosides and oxazolidinones, β-lactams are often used to treat nocardiosis. β-lactam resistance has however been reported for *Nocardia* with strong variations from one species to another. As stated in a retrospective study while about 80% of *Nocardia farcinica* are resistant to the β-lactam cefotaxime only 5% of the *Nocardia cyriacigeorgica* strains are (Lebeaux et al., 2019). β-lactamases are enzymes capable of hydrolyzing β-lactam antibiotics and are classified into four classes according to the Ambler nomenclature (Ambler et al., 1991; Bush, 2013). Classes A, C, and D are serine hydrolases while class B are metallo-β-lactamases that accommodate one or two zinc ions in their active site (Bahr et al., 2021). It is not yet fully documented if *Nocardia* species possess all four β-lactamases classes, however, they certainly widely harbor class A β-lactamases (ABL) (Valdezate et al., 2015). ABL are usually capable of hydrolyzing a wide range of β-lactams and have been found or can be found in the genomes of most *Nocardia* species. Only four studies have described the biochemical characterization of ABL, notably from *Nocardia lactamdurans* (Coque et al., 1993), *Nocardia farcinica* (Laurent et al., 1999; Lebeaux et al., 2019) and *Nocardia asteroides* (Poirel et al., 2001).

To date, there is no report of any three-dimensional structure of ABL from *Nocardia*. Such data could be helpful to fully apprehend the difference of substrate specificity and inhibitors efficacies observed between ABL from different *Nocardia* species (Laurent et al., 1999; Lebeaux et al., 2020; Poirel et al., 2001). To fill this gap, we describe in this study the biochemical and structural characterization of a ABL from *Nocardia cyriacigeorgica,* hereafter renamed NCY-1, one of the most encountered pathogenic human *Nocardia* species and for which ß-lactam resistance has been reported (Zhao et al., 2017).

## 2 Materials and methods

### 2.1 Gene amplification and cloning

Genomic DNA extracted from a *N. cyriacigeorgica* clinical isolate responsible for pulmonary nocardiosis was used as a template for polymerase chain reaction (PCR) with primers 5’- gatatgcaccacggcctgca-3’ and 5’-acggcgacgaagaagcgga-3’ by using Invitrogen™ Platinum SuperFi II DNA Polymerase (Thermo Fisher Scientific, Illkirch, France). A second PCR was performed on the aforementioned amplicon to amplify the truncated version of *NCY-1* with the following forward 5’-ggtaccgagaacctgtacttccagggttcggccgtggccgatccccggttcgccgcactggaaacg-3’and reverse 5’- gtggtgctcgagctaaccgagcacgtcgacgaccgtcctggtcgcgtcggc-3’primers. The purified PCR product was treated with DreamTaq polymerase (Thermo Fisher Scientific, Illkirch, France) to add overhanging A in 3’ and cloned into the Champion™ pET-SUMO (Invitrogen) following manufacturer instructions and the correct insertion of the DNA fragment was verified by sequencing.

### 2.2 Protein Expression and Purification

The pET-SUMO::*NCY-1* plasmid was transformed into the *E. coli* BL21 strain resistant to phage T1 (New England Biolabs) harboring the pRARE2 plasmid and plated onto LB+Agar plates supplemented with kanamycin (Kan) (50 μg.mL^-1^) and chloramphenicol (Cam) (30 μg.mL^-1^). An overnight (ON) preculture was used to inoculate 2 flasks containing each 3L of LB with Kan and Cam. When the cells reached the exponential phase, they were placed into ice for 40 min and protein expression was induced with 1 mM IPTG, and the culture was placed at 18°C ON. Cells were harvested by centrifugation for 15 min at 6000 *g* and resuspended in 80 mL of 50 mM Tris pH 8, 0.4 M NaCl, 5 mM β-mercaptoethanol, 1 mM benzamidine, 10 % glycerol, and 20 mM imidazole. Bacteria were lysed by sonication and centrifuged for 40 min at 27000 *g.* The clarified extract was loaded onto three gravity columns containing each 2 mL of Ni-NTA Sepharose beads (Cytiva) at 4°C. The columns were then washed with 25 mL of 50 mM Tris pH 8, 1 M NaCl, 5 mM β-mercaptoethanol, and 10% glycerol and the proteins were eluted with a total of 30 mL of buffer 50 mM Tris pH 8, 0.2 M NaCl, 5 mM β-mercaptoethanol, 10% glycerol, and 200 mM imidazole. At this stage about 230 mg of protein were recovered and the tag was cleaved by adding 4 mg of Tobacco Etch Virus (TEV) protease in a dialysis bag and dialyzed ON at 4°C against 50 mM Tris pH 8, 0.2 M NaCl, 5 mM β-mercaptoethanol and 10% glycerol. A second nickel affinity chromatography was performed to separate the cleaved NCY-1 from the tags and the Histagged-TEV protease. To do so, 2 columns with 3 mL beads each were equilibrated with the dialysis buffer and the samples were loaded three times. The columns were washed with 20 mL of dialysis buffer and 25 mL of 50 mM Tris pH 8, 1 M NaCl, 5 mM β-mercaptoethanol and 10% glycerol. About 200 mg of NCY-1 were recovered in the flow-through and the first wash step, concentrated to 5 mg.mL^-1^ and flash frozen in liquid nitrogen. For biochemical and crystallization purposes the protein was further purified by size-exclusion chromatography (SEC) on a Superdex 75 Increase 10/300 GL column (Cytiva) on which 5 mg of protein were injected and eluted with the following buffer: 20 mM Tris pH 8, 0.2 M NaCl, 0.5 mM β-mercaptoethanol and 5% glycerol.

### 2.3 Assessment of NCY-1 oligomeric state

The calibration curve was performed on a Superdex 75 Increase 10/300 GL column possessing a total volume (Vt) of 23.56 mL connected to an Äkta Go (Cytiva). The proteins were eluted in 20 mM Tris pH 8, 0.2 M NaCl and 0.5 mM β-mercaptoethanol. A mixture of 600 μg of bovine serum albumin, 200 μg of carbonic anhydrase from bovine erythrocytes, 190 μg of cytochrome c from horse heart and 400 μg of bovine aprotinin were injected simultaneously and eluted at a flow-rate of 0.3 mL.min^-1^. NCY-1 (300 μg) was injected independently and thyroglobulin (400 μg) was used to determine the void volume Vo. The volume of injection for each sample was 500 μL. The partition coefficient of each protein *K*_av_ was determined as follows: *K*_av_=(Vt-Vo)/(Ve-Vo) where Ve indicates the volume of elution of each protein. The calibration curve was obtained by plotting the *K*av against the Log of the molecular weight (M.W.)

### 2.4 Crystallization

Crystals were obtained in sitting drops using the Swissci MRC MAXI plate mixing 1μL of protein solution concentrated to 14 mg.mL^-1^ with 1μL of reservoir solution made of 0.1 M citric acid pH3.5, 10 % PEG 6000 and 6% ethylene glycol and equilibrated against a reservoir volume of 200 μL at 18°C. Crystals were cryoprotected by the addition of 4 μL of a solution made of 0.1 M citric acid pH 3.5, 14 % PEG 6000, and 20 % ethylene glycol, on top of the crystallization drop and incubated for 16 hours before being cryocooled in liquid nitrogen.

### 2.5 X-ray data collection, structure determination, and refinement

Data collection was performed on the PXI-X06SA beamline at the Swiss Light Source, Paul Scherrer Institute, Villigen, Switzerland. A total of 800 images were collected at an oscillation range of 0.2°, a crystal-to-detector distance of 145 mm, and a X-ray wavelength of 0.9999 Å. Data were processed and scaled with the XDS package (Kabsch, 2010). The phase problem was solved by molecular replacement using Phaser (McCoy, 2007) and the Phenix package (Liebschner et al., 2019), and an AlphaFold model (Jumper et al., 2021) of NCY-1 as a search template. The structure was further manually rebuilt with Coot (Casañal et al., 2020) and refined with the Phenix package (Liebschner et al., 2019). Statistical values given in Tables 3 and 4 were calculated with Aimless from the *CCP*4 suite (Agirre et al., 2023) and Phenix (Liebschner et al., 2019).

**Table 1.**
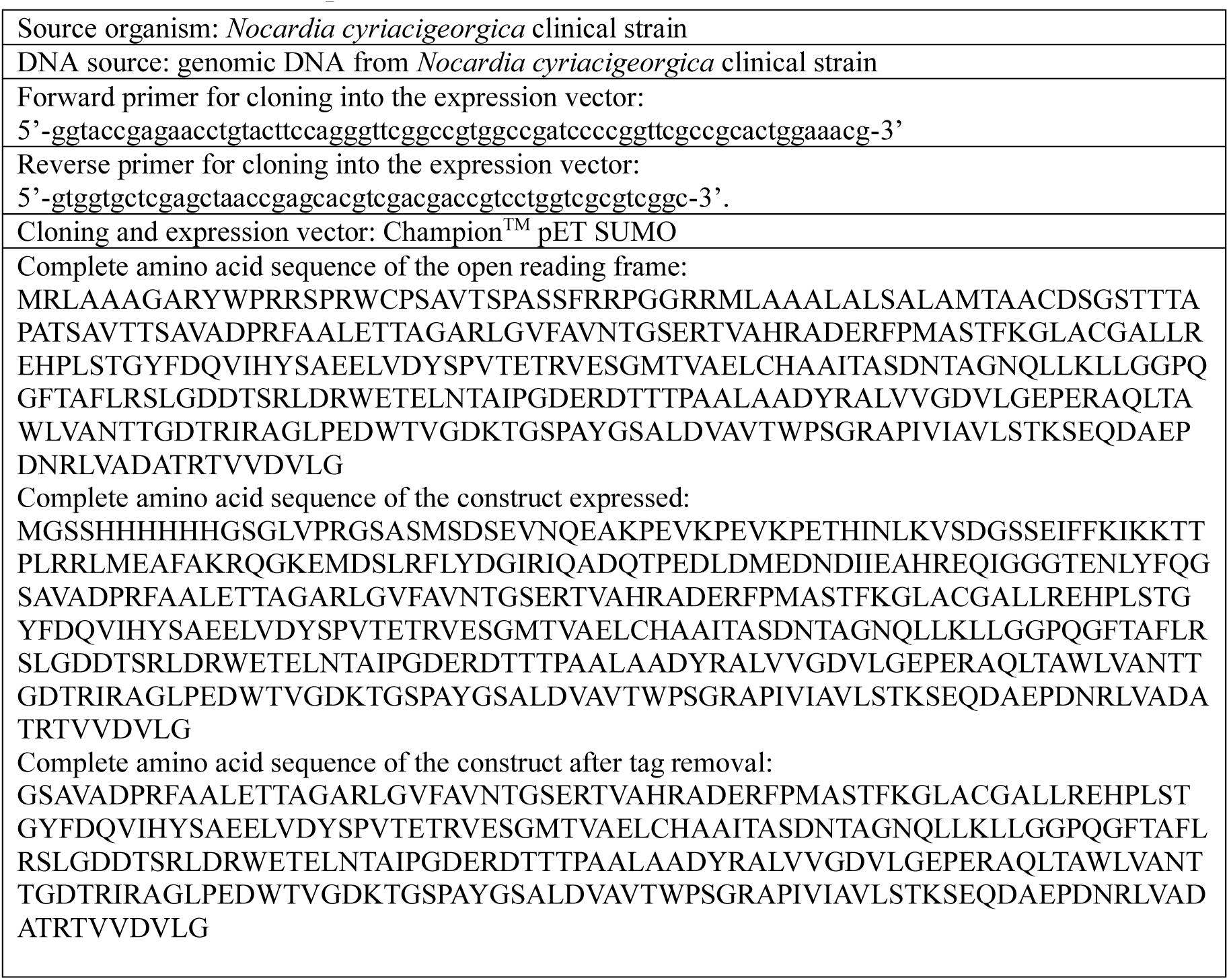
Macromolecule production information.

**Table 2.**
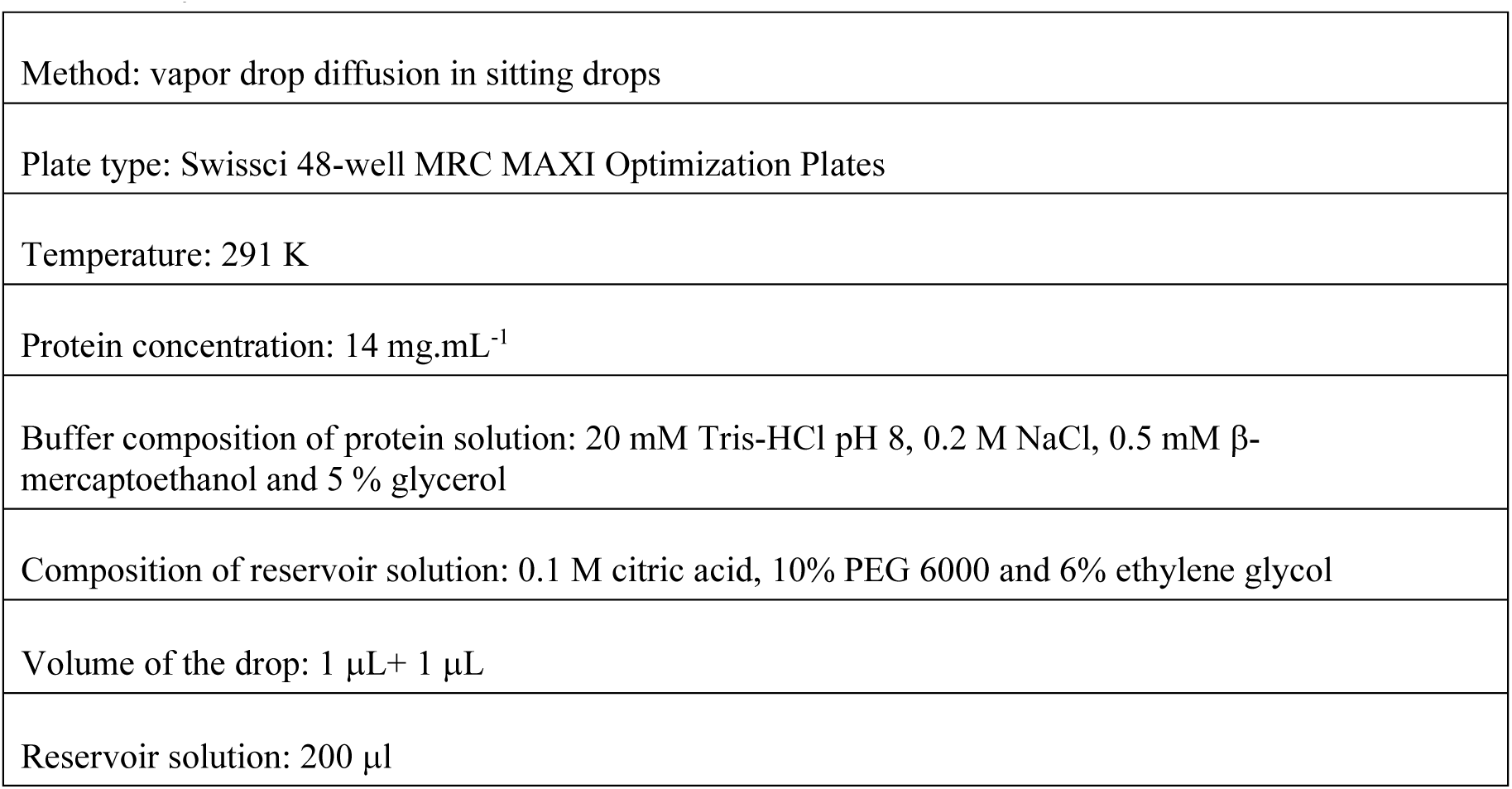
Crystallization.

|  |
| --- |
| Method: vapor drop diffusion in sitting drops |
| Plate type: Swissci 48-well MRC MAXI Optimization Plates |
| Temperature: 291 K |
| Protein concentration: 14 mg.mL<sup>-1</sup> |
| Buffer composition of protein solution: 20 mM Tris-HCl pH 8, 0.2 M NaCl, 0.5 mM β-mercaptoethanol and 5 % glycerol |
| Composition of reservoir solution: 0.1 M citric acid, 10% PEG 6000 and 6% ethylene glycol |
| Volume of the drop: 1 μL + 1 μL |
| Reservoir solution: 200 μl |

**Table 3.** Data collection statistics.

|  |  |
| --- | --- |
| **Beamline** | PXI-X06SA SLS |
| **Wavelength** | 0.9999 |
| **Data collection temperature (K)** | 100 |
| **Resolution range** | 43.95 – 1.25 (1.28 – 1.25) |
| **Space group** | $P 3_2 2 1$ |
| **Unit cell ($\text{\AA}$,$^\circ$)** | a=b=54.18, c=131.86, $\alpha$=$\beta$=90, $\gamma$=120 |
| **Solvent content (%)** | 38.48 |
| **Total reflections** | 542218 (27178) |
| **Unique reflections** | 63063 (3101) |
| **Multiplicity** | 8.6 (8.8) |
| **Completeness (%)** | 100 (100) |
| **Mean I/sigma(I)** | 12 (1) |
| **Wilson B-factor ($\text{\AA}^2$)** | 17.7 |
| **R-meas (%)** | 7.8 (238.6) |
| **CC1/2** | 0.99 (0.28) |

Statistics for the highest-resolution shell are shown in parentheses.

**Table 4.**
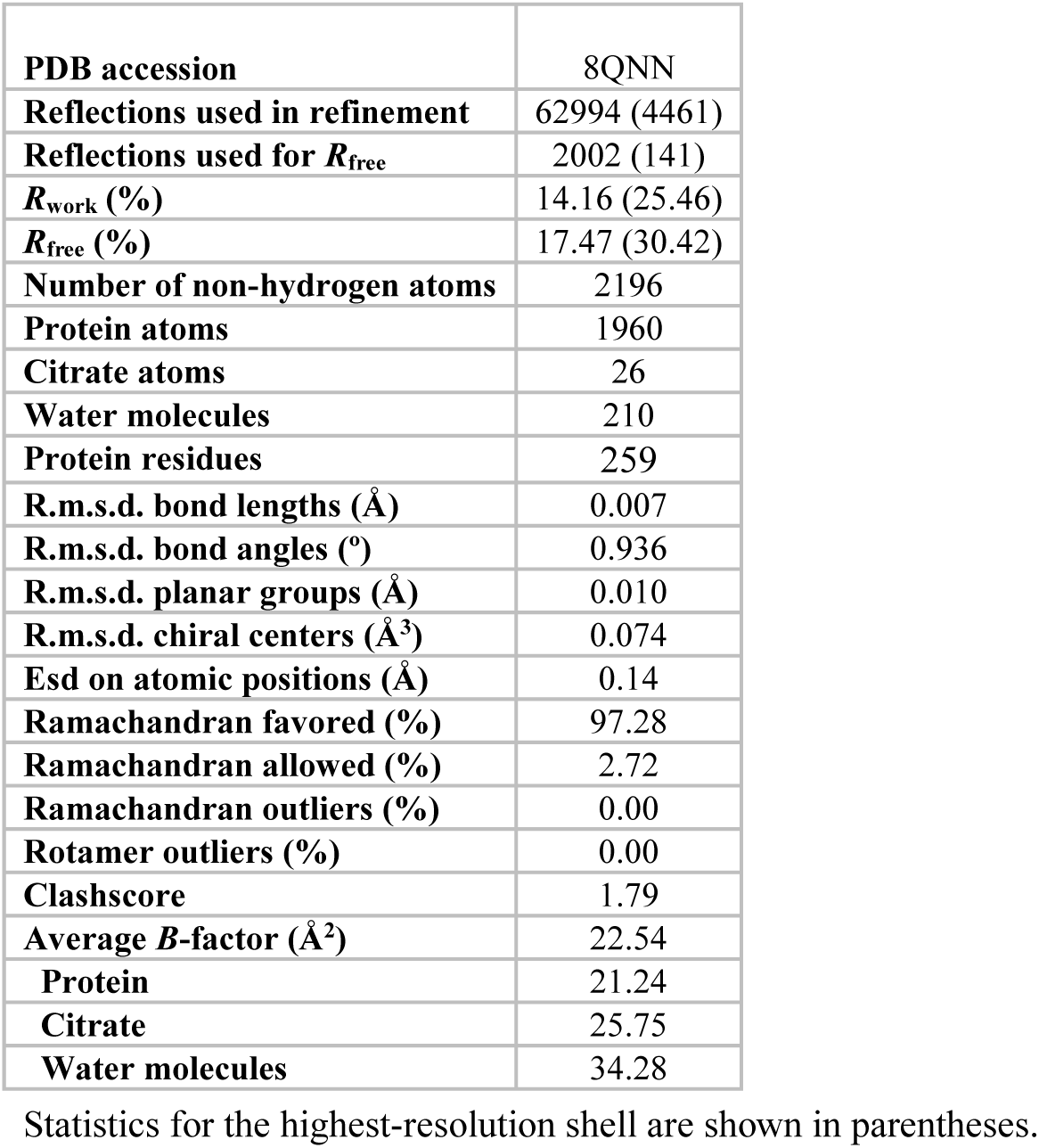
Refinement statistics.

### 2.6 Activity assays

The β-lactamase assay was performed in 96-well plates and a reaction volume of 100 μL in 1x phosphate saline (PBS) buffer pH 7 and the presence of 0.2 mM nitrocefin. Reactions were initiated by addition of 2, 4, or 8 nM of enzyme or 8 nM of enzyme incubated with 500 μM of avibactam for 30 min. A control reaction was performed with all components but the enzyme. The optical density at 486 nm was recorded for 75 minutes with data points acquired every 1 min on a Tecan Infinite^®^ 200 PRO M Plex plate reader.

## 3 Results and discussion

### 3.1 Gene amplification, construct design and protein expression

We first tried to amplify a ß-lactamase gene from the *N. cyriacigeorgica* clinical strain using primers based on the sequence of *Nocardia cyriacigeorgica* GUH-2 chromosome complete genome GenBank: FO082843.1. The set of primers led to the amplification of a DNA fragment encompassing the ß-lactamase AST-1 precursor GenBank:CCF63029.1. We then analyzed the putative open reading frame resulting from this nucleotide sequence and blast it to assess the sequence conservation. The final nucleotide sequence was deposited at the NCBI, GenBank under the following accession number OR515607.

The translation of the nucleotide indicates an open reading frame (ORF) of 333 amino acids. As ABL are synthesized as a precursor in the cytoplasm and then secreted into the periplasm (Kaderabkova et al., 2022), we explored the possibility of the presence of a peptide signal. We analyzed the sequence with the Signal P server (Teufel et al., 2022) which enabled us to identify a probable secretion signal with a cleavage site between residues Ala53 and Cys54. To further design an optimal sequence for protein expression and crystallization, we made a predictive three-dimensional model with AlphaFold (Jumper et al., 2021) to identify flexible/non-folded regions. This last analysis prompted us to design an ORF ranging from residues Ser71 to Gly333.

Finally, following this design approach, we cloned the nucleotide sequence of NCY-1 into an expression vector enabling fusion with the SUMO tag in the N-terminus as well as a multi-histidine tag. We also added between the tag and the coding sequence a sequence encoding for the Tobacco Etch Virus protease to remove the tag during the purification process. This strategy led to the production of a highly soluble form of NCY-1, and following a two-step purification procedure, we could purify about 200 mg of protein from 6 liters of culture with two nickel affinity chromatography steps. The protein was further purified on SEC leading to a very pure and homogeneous protein as shown in **Figure 1(*a*)**.

**Figure 1.**
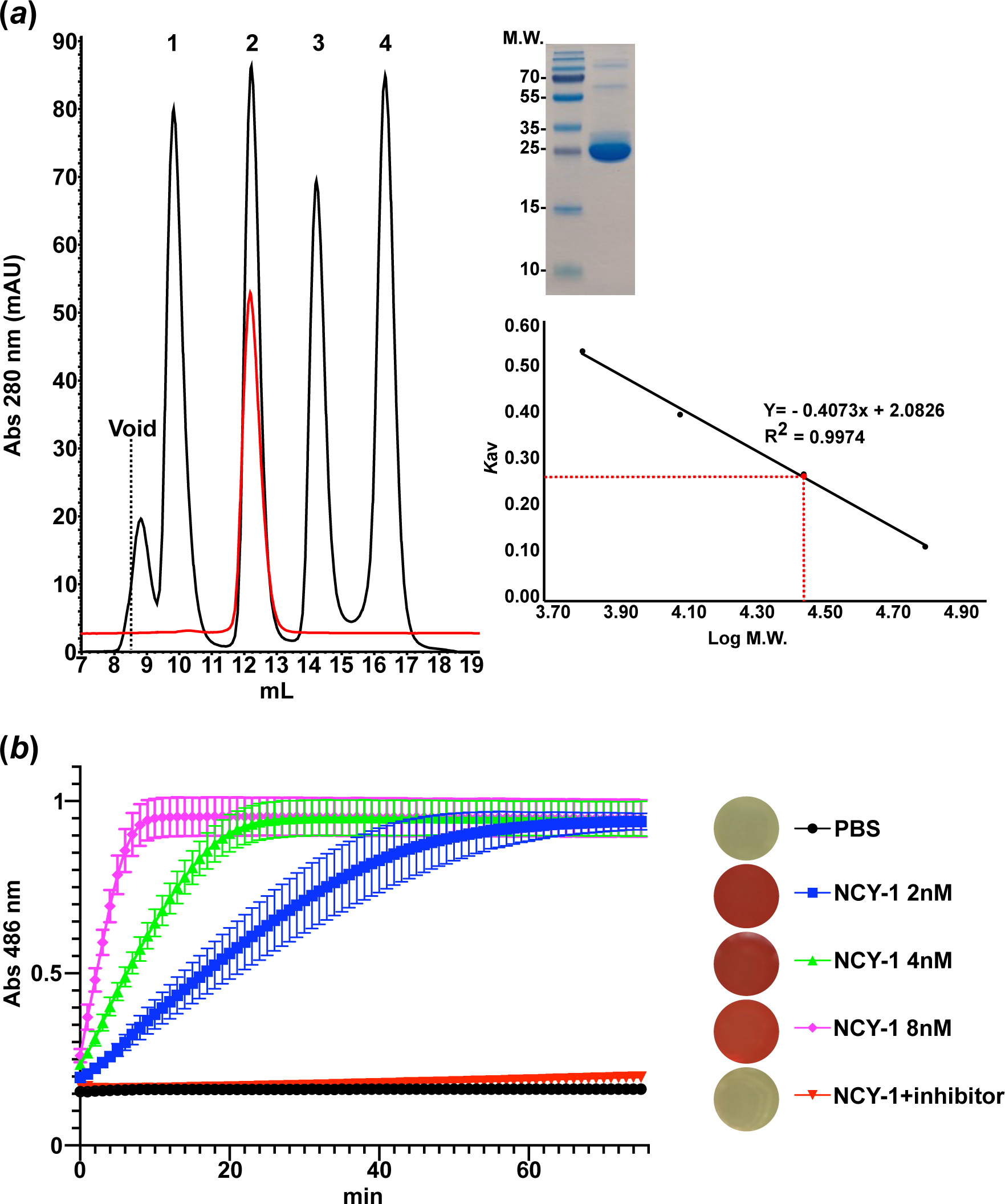
Evaluation of the protein quality. (***a***) Size-exclusion chromatography profile of NCY-1 on Superdex 75 Increase 10/300 GL column. The elution profile was compared to protein standard: bovine serum albumin (66 kDa, Ve=9.82 mL), bovine carbonic anhydrase (29 kDa, Ve=12.22 mL), horse cytochrome c (12.9 kDa, ve=14.22) and bovine aprotinin (6.5 kDa, Ve=16.33 mL). Ve indicates the elution volume. NCY-1 was eluted at a volume of 12.18 mL. The void volume (Vo) was determined at 8.53 mL with the volume of elution of thyroglobulin (M.W. 669kDa). The NCY-1 (10 μg loaded) sample used for biochemical and structural analysis is shown on the 15 % Coomassie stained denaturing gel. **(*b*)** Determination of the β-lactamase activity of NCY-1 in the presence of nitrocefin. Three concentrations of the enzyme were tested (2 nM, blue curve; 4 nM, green curve; 8 nM, pink curve) and two control experiments were performed: the reaction background (black curve) is the reaction done with PBS instead of enzyme and one with avibactam (red curve), a potent covalent inhibitor of β-lactamases. The data from three technical replicates were analyzed with GraphPad Prism version 10.0.2. On the right of the curves are shown pictures taken at the end of the reaction for each well and attesting to the color shift from yellow to red upon enzymatic reaction.

### 3.2 Protein assessment quality

To ensure our construct design strategy was successful and led to a well-folded and functional protein, we first assessed the behavior of the protein on SEC. The calibration curve attested that the NCY-1 behaves as a homogeneous and stable monomer of an apparent molecular weight (M.W.) of about 29 kDa and as expected from the theoretical M.W. and for this class of enzyme known to be a monomer **(Figure 1(*a*))**. We also conducted a ß-lactamase activity assay. To do so, we used nitrocefin a chromogenic cephalosporin substrate that shifts from yellow to red color upon ß-lactamase activity. The presence of nM concentrations of pure NCY-1 triggered a color shift as shown in **Figure 1(*b*)**. The absorbance monitoring assay attested of an increased absorbance at 486 nm in a NCY-1 concentration-dependent manner and in contrast to the two control experiments performed without enzyme or with enzyme incubated with avibactam, a covalent inhibitor of β-lactamases. Altogether, these data are in agreement that the construct design was successful and led to the production of a functional protein.

### 3.3 Crystallization, data collection, and structure determination

We initiated the first crystallization trial at two different protein concentrations of 5 and 20.9 mg.mL^-1^. Most drops were clear indicating the very soluble behavior of the protein. Nonetheless, several conditions have promoted the appearance of NCY-1 crystals. The best crystals were optimized in sitting drops at a protein concentration of 14 mg.mL^-1^ and using a reservoir solution made of 0.1 M citric acid pH 3.5, 10 % PEG 6000, and 6 % ethylene glycol.

A full X-ray dataset was collected and processed to a resolution of 1.25Å **(Table 3)**. The structure was solved by molecular replacement using the prediction model mentioned above as a search request. We finally rebuilt and refined the model to R_work_/R_free_ values of 14.16%/17.47% with good geometry **(Table 4)**. The asymmetric unit possesses one monomer. Most of the residues could be rebuilt except residues 71 to 73 and the G in the N-terminus from the TEV cleavage site. The last G333 in the C-terminus could not be seen as well. In addition, we unambiguously identified two molecules of citrate that was present in the crystallization condition. One of the citrates is situated on the symmetry axis on the surface of the protein and contributes to the crystal packing of symmetry related NCY-1 monomers and through interactions with residues A164, E165, H168, Q250 and W254 (not shown). The second citrate is found in the active site and its interaction is further described below.

### 3.4 Structural analysis

NCY-1 has a classical ABL structure with an α/β hydrolase fold made of two domains **(Figure 2(*a*))**. The first domain is a mixed α/β fold, made of strands β1 to β5 and helices α1 and α9. The second domain is fully α-helical formed by helices 2, 3, 4, 5, 6, 7 and 8 **(Figure 2(*a*))**. The structure is made of nine α-helices and six helices of 3_10_ type. Five β-strands (β1-5) form one antiparallel β-sheet and complete the structure. For clarity and to match the standard nomenclature of ABL the residue indicated between parentheses are following the Ambler numbering scheme for class A (Ambler et al., 1991).

**Figure 2.**
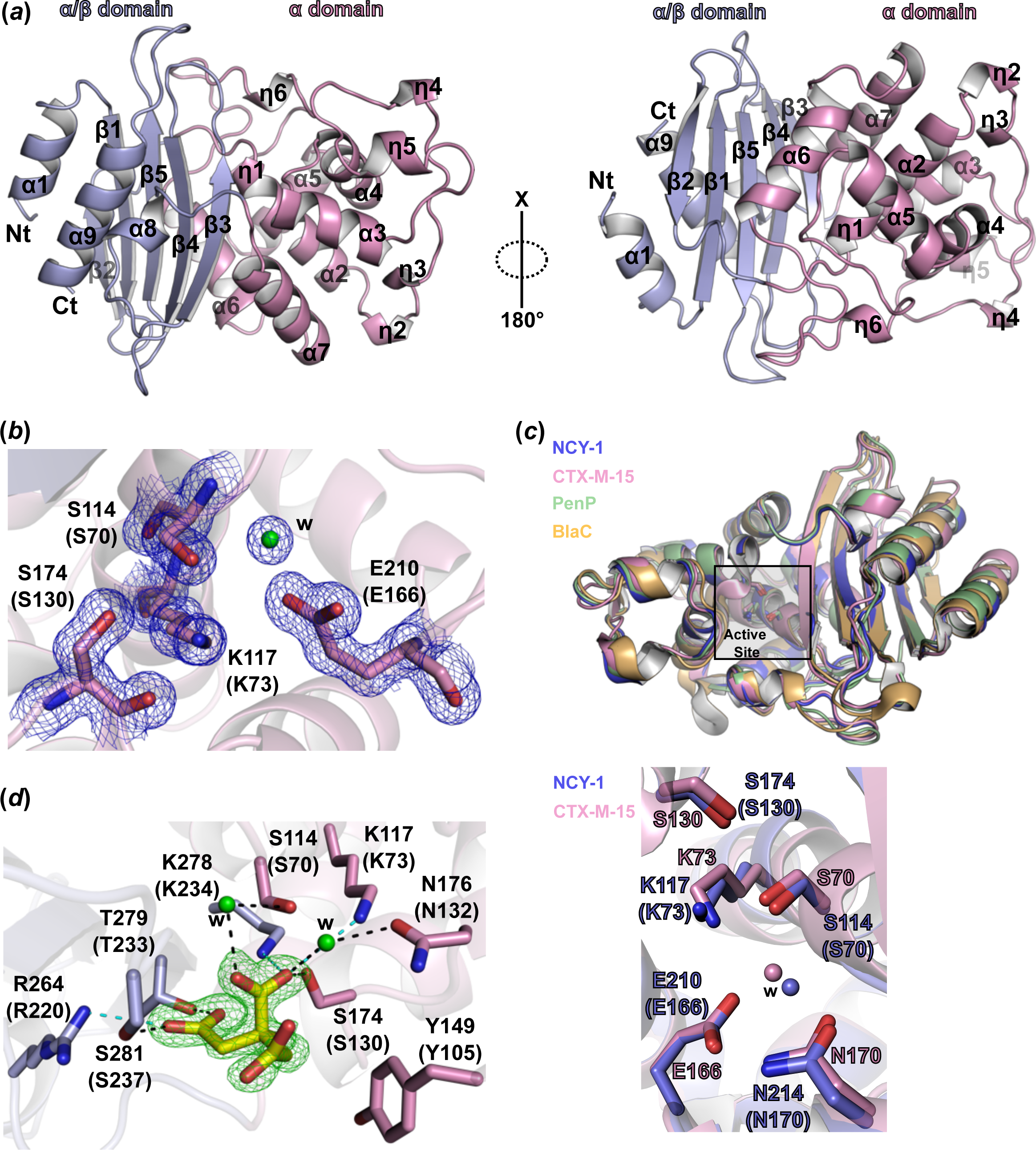
Structural analysis of NCY-1. (***a***) Cartoon representation of the NCY-1 three-dimensional structure in two different orientations. The α/β domain is in blue and the α domain in pink. Nt and Ct stand for N-terminus and C-terminus respectively. **(*b*)** Depiction of the important residues for catalysis of class A β-lactamase activity. The 2Fo-Fc map (blue mesh) contoured at a level of 1 σ attests to the high quality of the electron density map. W indicates water molecules that are displayed as a green sphere. Residues in parentheses indicate Ambler numbering. **(*c*)** Structural comparison of NCY-1 (blue) with its closest homologues ABL : PenP from *Bacillus licheniformis* (green), CTX-M-15 (pink) and BlaC from *Mycobacterium tuberculosis* (orange). The panel below displays a close-up of the active site of CTX-M-15 (pink) and NCY-1 (blue) and attests to the strict conservation of catalytically important residues, shown as sticks and water molecules (w). **(*d*)** Network of interactions between the citrate molecule and NCY-1 residues. The dashed black and cyan lines indicate hydrogen bonds and salt-bridges respectively. Nitrogen atoms are colored in blue and oxygen in red. The interactions were analyzed with the PLIP server (Adasme et al., 2021). The simulated annealing omit map proving the presence of the citrate, is shown as a yellow stick. The map is contoured at a level of 3.5 σ and represented as a green mesh.

On the contrary to some other class A, we did not notice any disulfide bridge albeit the C121 (C77) and C167 (C123) side chains on helices α2 and α3 are in close proximity. The absence of a disulfide bridge could be the result of the recombinant expression of the protein in the cytoplasm of *E. coli* as we removed the secretion signal of NCY-1. The cytoplasm is indeed less favorable for the formation of disulfide bonds than the periplasm where oxidative conditions are met and where β-lactamases are normally located.

The active site residues and water molecules in proximity are well-ordered and well-defined in the electron density map **(Figure 2(*b*))**. The catalytic amino acids are at the interface of the two domains and are composed of residues S114 (S70), K117 (K73), Ser174 (S130), and Glu210 (E166) **(Figure 2(*b*))**. These amino acids are strictly conserved when compared with other ABL structures from other phylae and particularly the well-characterized CTX-M-15 **(Figure 2(*c*) and 3)** and for which these residues were shown to be crucial for the hydrolysis of β-lactam moiety (Tooke et al., 2019).

**Figure 3.**
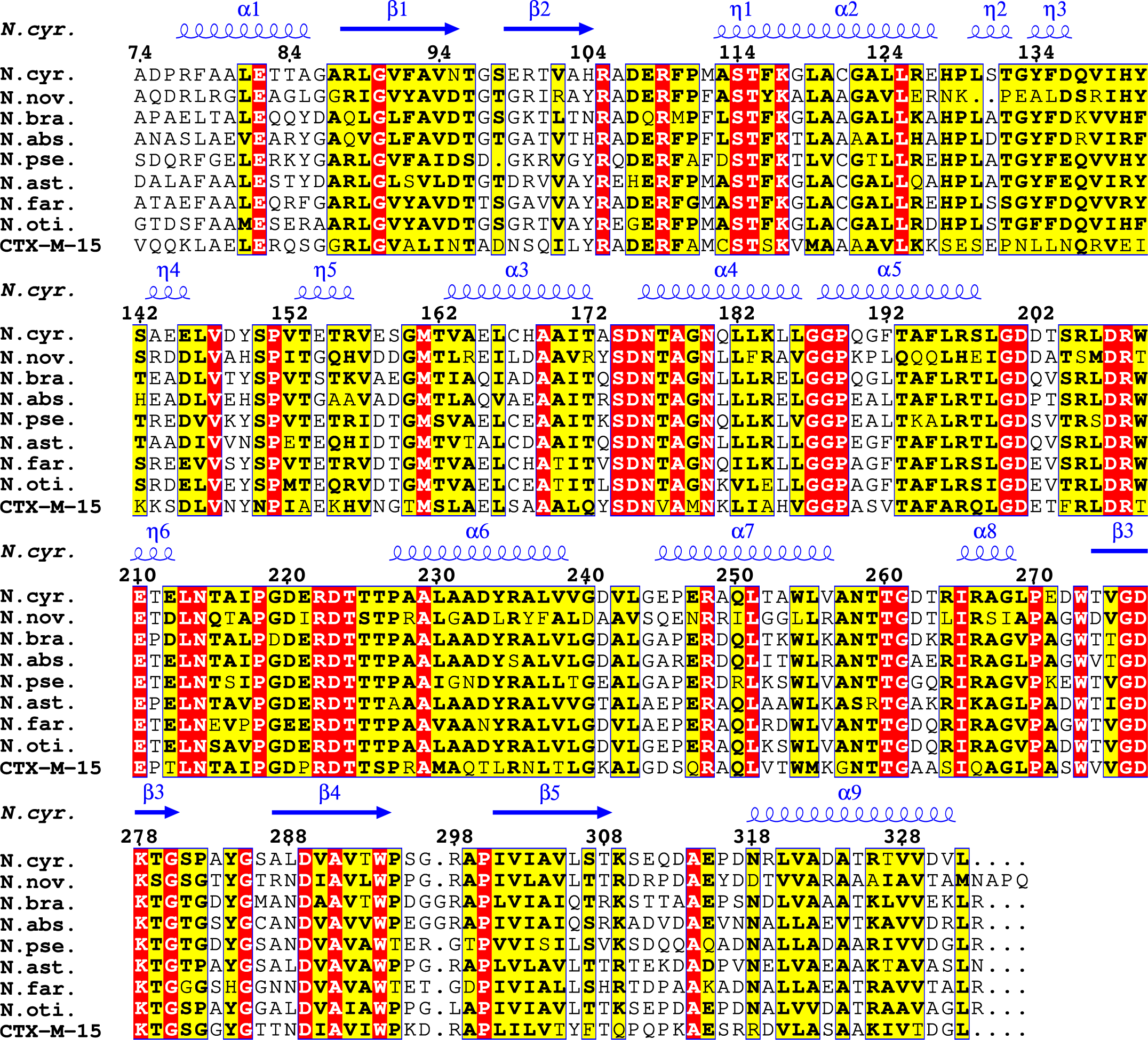
Multiple sequence alignment of class A β-lactamases from human pathogenic *Nocardia*. The sequence of NCY-1 (*N. cyr.*) was aligned with the sequences of ABL from *Nocardia otitidiscaviarum* NEB252 (N. oti.), *Nocardia farcinica* IFM 10152 (N. far.), *Nocardia brasiliensis* ATCC 700358 (N. bra.), *Nocardia asteroides* NCTC11293 (N. ast.), *Nocardia nova* SH22a (N. nov.), *Nocardia pseudobrasiliensis* ATCC 51512 (N.pse.), *Nocardia abscessus* NBRC 100374 (N. abs.) and CTX-M-15. The alignment displays only the sequences matching the NCY-1 residues that were modeled in the structure; thus, some of the sequences are missing some residues in the N-terminus. The secondary structure elements from the NCY-1 structure are placed above the sequence alignment where α, β and, η indicates α-helix, β-sheet and η, 3_10_ helix respectively. The figure was generated with the ENDscript server (Robert and Gouet, 2014) https://endscript.ibcp.fr.

Further analysis shows that the ABL sequence signature (Philippon et al., 2019) is conserved notably in the three motifs 114-STFK-117 (70-STFK-73), 174-SDN-176 (130-SDN-132), and 278-KTG-280 (234-KTG-236) **(Figure 3)**. The E210 (E166) residue on the so-called Ω-loop, and important for ABL enzymatic activity is also conserved. A detailed classification of β-lactamases based on multiple sequence alignment attested that subclass A1 (Philippon et al., 2019) possesses a clear signature sequence that presents on top of the three motifs mentioned above eighteen additional conserved residues. NCY-1 harbors strict conservation of these consensus residues **(Figure 3)** as listed further: R105 (61), R109 (65), T115 (71), G122 (78), A169 (125), N180 (136), D201 (157), R208 (164), N214 (170), D223 (179), T224 (180), T225 (181), T226 (182), P227 (183), W254 (210), R266 (222), W273 (229) and D277 (233).

We searched for the closest structures of NCY-1 in the entire protein database using the PDBeFold server (https://www.ebi.ac.uk/msd-srv/ssm/ssmstart.html) (Krissinel and Henrick, 2004). Evidently, because of the high number of ABL whose structures were determined, we retrieved numerous similar structures with RMSD below 1 Å. The closest structures were the ABL from *Bacillus licheniformis* (PDB code 1i2s) and named PenP with a RMSD value of 0.7 Å and 45 % protein sequence identity and more interestingly close similarity (RMSD of 0.78 Å and 46 % protein sequence identity) with the extended-spectrum β-lactamase CTX-M-15 from *Escherichia coli* (PDB code 4hbt) (Lahiri et al., 2013) **(Figure 2(*c*))**. We found that ABL from species of the same order as *Nocardia*, namely BlaC from *Mycobacterium tuberculosis* (PDB code 2gdn) (Wang et al., 2006) is slightly more distant as it shares a RMSD of 1 Å and 50 % sequence identity. Further, a comparison of the active site of NCY-1 with the canonical CTX-M-15 demonstrates strict conservation of key and important amino acids for the catalysis as well as the same positioning of the water molecule crucial for the hydrolysis of the β-lactam ring **(Figure 2(*c*))**. This analysis suggests therefore that NCY-1 might have a broad-spectrum substrate specificity as its closest homologues.

Additionally, we could unambiguously identify one citrate molecule that is tightly bound in the vicinity of the active site and as supported by the simulated-annealing Fo-Fc omit map, **(Figure 2(*d*))**. The side chain of Y149 (Y105) interacts through van der Waals interactions **(Figure 2(*d*))**. A network of H-bonds between the side chains of S114 (S70), S174 (S130), T279 (T235), and S281(S237) allows interaction with the citrate molecule, while residues S114 (S70) and N176 (N132) via water molecules mediate specific interactions. The tight binding of citrate is achieved by salt bridges involving K117 (K73), R264 (R220), and K278 (K234) **(Figure 2(*d*))**. A similar interaction with citrate was previously observed for the structure of the TEM-72, a class A β-lactamase from *Enterobacteriaceae* (Docquier et al., 2011).

Finally, to address whether the NCY-1 structure is representative of ABL from other *Nocardia* species, we performed multiple sequence alignments of ABL from some of the most frequently encountered human pathogenic *Nocardia* spp. **(Figure 3)**. This sequence analysis demonstrates a close conservation and sequence identity of NCY-1 with the *N. nova* 49 %, *N. brasiliensis* 66 %, *N. abscessus* 65 %, *N. pseudobrasiliensis* 65 %, *N. asteroides* 69 %, *N. farcinica* 72 % and *N. otitidiscaviarum* 77 %, and 45% with CTX-M-15. All the proteins from *Nocardia* spp. have almost a strict conservation of the three motifs described earlier and the eighteen residues characterizing class A1. Only a few substitutions were noticed for the class A1 signature in *N. abscessus*, *N. asteroides*, and *N. nova* but probably with no consequence for the enzyme function as they do not concern residues critical for the activity of ABL (Tooke et al., 2019). Thus, this analysis strongly suggests that all ABL from pathogenic *Nocardia* species have very likely similar structures and that all these species may possess functional ABL, as already shown for *N. farcinica* and *N. asteroides* (Laurent et al., 1999; Poirel et al., 2001).

## Acknowledgements

We thank the CNRS, the IRIM directory and Dr. K. Brodolin for support. We acknowledge the Paul Scherrer Institute, Villigen, Switzerland for the provision of synchrotron radiation beamtime at beamline PXI-X06SA of the SLS. We thank Dr. B. Iorga for the fruitful discussions.

## Funding information

JC PhD fellowship is financed by the CNRS.

## Notes

### Competing Interest Statement

The authors have declared no competing interest.

### Summary of Updates

Some parts of the text have been clarified, Table 3 and 4 have been updated. Figure 2 and Figure 3 have been modified.

## Bibliography

Adasme, M.F., Linnemann, K.L., Bolz, S.N., Kaiser, F., Salentin, S., Haupt, V.J., Schroeder, M., 2021. PLIP 2021: expanding the scope of the protein–ligand interaction profiler to DNA and RNA. Nucleic Acids Res. 49, W530–W534. 10.1093/nar/gkab294

Agirre, J., Atanasova, M., Bagdonas, H., Ballard, C.B., Baslé, A., Beilsten-Edmands, J., Borges, R.J., Brown, D.G., Burgos-Mármol, J.J., Berrisford, J.M., Bond, P.S., Caballero, I., Catapano, L., Chojnowski, G., Cook, A.G., Cowtan, K.D., Croll, T.I., Debreczeni, J.É., Devenish, N.E., Dodson, E.J., Drevon, T.R., Emsley, P., Evans, G., Evans, P.R., Fando, M., Foadi, J., Fuentes-Montero, L., Garman, E.F., Gerstel, M., Gildea, R.J., Hatti, K., Hekkelman, M.L., Heuser, P., Hoh, S.W., Hough, M.A., Jenkins, H.T., Jiménez, E., Joosten, R.P., Keegan, R.M., Keep, N., Krissinel, E.B., Kolenko, P., Kovalevskiy, O., Lamzin, V.S., Lawson, D.M., Lebedev, A.A., Leslie, A.G.W., Lohkamp, B., Long, F., Malý, M., McCoy, A.J., McNicholas, S.J., Medina, A., Millán, C., Murray, J.W., Murshudov, G.N., Nicholls, R.A., Noble, M.E.M., Oeffner, R., Pannu, N.S., Parkhurst, J.M., Pearce, N., Pereira, J., Perrakis, A., Powell, H.R., Read, R.J., Rigden, D.J., Rochira, W., Sammito, M., Sánchez Rodríguez, F., Sheldrick, G.M., Shelley, K.L., Simkovic, F., Simpkin, A.J., Skubak, P., Sobolev, E., Steiner, R.A., Stevenson, K., Tews, I., Thomas, J.M.H., Thorn, A., Valls, J.T., Uski, V., Usón, I., Vagin, A., Velankar, S., Vollmar, M., Walden, H., Waterman, D., Wilson, K.S., Winn, M.D., Winter, G., Wojdyr, M., Yamashita, K., 2023. The CCP4 suite: integrative software for macromolecular crystallography. Acta Crystallogr. Sect. Struct. Biol. 79, 449–461. 10.1107/S2059798323003595

Ambler, R.P., Coulson, A.F., Frère, J.M., Ghuysen, J.M., Joris, B., Forsman, M., Levesque, R.C., Tiraby, G., Waley, S.G., 1991. A standard numbering scheme for the class A beta-lactamases. Biochem. J. 276 ( Pt 1), 269–270. 10.1042/bj2760269

Bahr, G., González, L.J., Vila, A.J., 2021. Metallo-β-lactamases in the age of multidrug resistance: from structure and mechanism to evolution, dissemination and inhibitor design. Chem. Rev. 121, 7957–8094. 10.1021/acs.chemrev.1c00138

Bush, K., 2013. The ABCD’s of β-lactamase nomenclature. J. Infect. Chemother. Off. J. Jpn. Soc. Chemother. 19, 549–559. 10.1007/s10156-013-0640-7

Casañal, A., Lohkamp, B., Emsley, P., 2020. Current developments in Coot for macromolecular model building of Electron Cryo-microscopy and Crystallographic Data. Protein Sci. Publ. Protein Soc. 29, 1069–1078. 10.1002/pro.3791

Cattaneo, C., Antoniazzi, F., Caira, M., Castagnola, C., Delia, M., Tumbarello, M., Rossi, G., Pagano, L., 2013. Nocardia spp infections among hematological patients: results of a retrospective multicenter study. Int. J. Infect. Dis. IJID Off. Publ. Int. Soc. Infect. Dis. 17, e610–614. 10.1016/j.ijid.2013.01.013

Coque, J.J., Liras, P., Martín, J.F., 1993. Genes for a beta-lactamase, a penicillin-binding protein and a transmembrane protein are clustered with the cephamycin biosynthetic genes in Nocardia lactamdurans. EMBO J. 12, 631–639. 10.1002/j.1460-2075.1993.tb05696.x

Docquier, J.-D., Benvenuti, M., Calderone, V., Rossolini, G.-M., Mangani, S., 2011. Structure of the extended-spectrum β-lactamase TEM-72 inhibited by citrate. Acta Crystallograph. Sect. F Struct. Biol. Cryst. Commun. 67, 303–306. 10.1107/S1744309110054680

Hershko, Y., Levytskyi, K., Rannon, E., Assous, M.V., Ken-Dror, S., Amit, S., Ben-Zvi, H., Sagi, O., Schwartz, O., Sorek, N., Szwarcwort, M., Barkan, D., Burstein, D., Adler, A., 2023. Phenotypic and genotypic analysis of antimicrobial resistance in Nocardia species. J. Antimicrob. Chemother. dkad236. 10.1093/jac/dkad236

Hodille, E., Prudhomme, C., Dumitrescu, O., Benito, Y., Dauwalder, O., Lina, G., 2023. Rapid, Easy, and Reliable Identification of Nocardia sp. by MALDI-TOF Mass Spectrometry, VITEK®-MS IVD V3.2 Database, Using Direct Deposit. Int. J. Mol. Sci. 24, 5469. 10.3390/ijms24065469

Jumper, J., Evans, R., Pritzel, A., Green, T., Figurnov, M., Ronneberger, O., Tunyasuvunakool, K., Bates, R., Žídek, A., Potapenko, A., Bridgland, A., Meyer, C., Kohl, S.A.A., Ballard, A.J., Cowie, A., Romera-Paredes, B., Nikolov, S., Jain, R., Adler, J., Back, T., Petersen, S., Reiman, D., Clancy, E., Zielinski, M., Steinegger, M., Pacholska, M., Berghammer, T., Bodenstein, S., Silver, D., Vinyals, O., Senior, A.W., Kavukcuoglu, K., Kohli, P., Hassabis, D., 2021. Highly accurate protein structure prediction with AlphaFold. Nature 596, 583–589. 10.1038/s41586-021-03819-2

Kabsch, W., 2010. XDS. Acta Crystallogr. D Biol. Crystallogr. 66, 125–132. 10.1107/S0907444909047337

Kaderabkova, N., Bharathwaj, M., Furniss, R.C.D., Gonzalez, D., Palmer, T., Mavridou, D.A.I., 2022. The biogenesis of β-lactamase enzymes. Microbiology 168. 10.1099/mic.0.001217

Krissinel, E., Henrick, K., 2004. Secondary-structure matching (SSM), a new tool for fast protein structure alignment in three dimensions. Acta Crystallogr. D Biol. Crystallogr. 60, 2256– 2268. 10.1107/S0907444904026460

Lahiri, S.D., Mangani, S., Durand-Reville, T., Benvenuti, M., De Luca, F., Sanyal, G., Docquier, J.-D., 2013. Structural Insight into Potent Broad-Spectrum Inhibition with Reversible Recyclization Mechanism: Avibactam in Complex with CTX-M-15 and Pseudomonas aeruginosa AmpC β-Lactamases. Antimicrob. Agents Chemother. 57, 2496–2505. 10.1128/aac.02247-12

Laurent, F., Poirel, L., Naas, T., Chaibi, E.B., Labia, R., Boiron, P., Nordmann, P., 1999. Biochemical-genetic analysis and distribution of FAR-1, a class A beta-lactamase from Nocardia farcinica. Antimicrob. Agents Chemother. 43, 1644–1650. 10.1128/AAC.43.7.1644

Lebeaux, D., Bergeron, E., Berthet, J., Djadi-Prat, J., Mouniée, D., Boiron, P., Lortholary, O., Rodriguez-Nava, V., 2019. Antibiotic susceptibility testing and species identification of Nocardia isolates: a retrospective analysis of data from a French expert laboratory, 2010-2015. Clin. Microbiol. Infect. Off. Publ. Eur. Soc. Clin. Microbiol. Infect. Dis. 25, 489–495. 10.1016/j.cmi.2018.06.013

Lebeaux, D., Ourghanlian, C., Dorchène, D., Soroka, D., Edoo, Z., Compain, F., Arthur, M., 2020. Inhibition Activity of Avibactam against Nocardia farcinica β-Lactamase FARIFM10152. Antimicrob. Agents Chemother. 64, e01551–19. 10.1128/AAC.01551-19

Liebschner, D., Afonine, P.V., Baker, M.L., Bunkóczi, G., Chen, V.B., Croll, T.I., Hintze, B., Hung, L.-W., Jain, S., McCoy, A.J., Moriarty, N.W., Oeffner, R.D., Poon, B.K., Prisant, M.G., Read, R.J., Richardson, J.S., Richardson, D.C., Sammito, M.D., Sobolev, O.V., Stockwell, D.H., Terwilliger, T.C., Urzhumtsev, A.G., Videau, L.L., Williams, C.J., Adams, P.D., 2019. Macromolecular structure determination using X-rays, neutrons and electrons: recent developments in Phenix. Acta Crystallogr. Sect. Struct. Biol. 75, 861–877. 10.1107/S2059798319011471

Margalit, I., Lebeaux, D., Tishler, O., Goldberg, E., Bishara, J., Yahav, D., Coussement, J., 2021. How do I manage nocardiosis? Clin. Microbiol. Infect. Off. Publ. Eur. Soc. Clin. Microbiol. Infect. Dis. 27, 550–558. 10.1016/j.cmi.2020.12.019

McCoy, A.J., 2007. Solving structures of protein complexes by molecular replacement with Phaser. Acta Crystallogr. D Biol. Crystallogr. 63, 32–41. 10.1107/S0907444906045975

McNeil, M.M., Brown, J.M., 1994. The medically important aerobic actinomycetes: epidemiology and microbiology. Clin. Microbiol. Rev. 7, 357–417. 10.1128/CMR.7.3.357

Mehta, H.H., Shamoo, Y., 2020. Pathogenic Nocardia: A diverse genus of emerging pathogens or just poorly recognized? PLoS Pathog. 16, e1008280. 10.1371/journal.ppat.1008280

Philippon, A., Jacquier, H., Ruppé, E., Labia, R., 2019. Structure-based classification of class A beta-lactamases, an update. Curr. Res. Transl. Med. 67, 115–122. 10.1016/j.retram.2019.05.003

Poirel, L., Laurent, F., Naas, T., Labia, R., Boiron, P., Nordmann, P., 2001. Molecular and biochemical analysis of AST-1, a class A beta-lactamase from Nocardia asteroides sensu stricto. Antimicrob. Agents Chemother. 45, 878–882. 10.1128/AAC.45.3.878-882.2001

Robert, X., Gouet, P., 2014. Deciphering key features in protein structures with the new ENDscript server. Nucleic Acids Res. 42, W320–W324. 10.1093/nar/gku316

Teufel, F., Almagro Armenteros, J.J., Johansen, A.R., Gíslason, M.H., Pihl, S.I., Tsirigos, K.D., Winther, O., Brunak, S., von Heijne, G., Nielsen, H., 2022. SignalP 6.0 predicts all five types of signal peptides using protein language models. Nat. Biotechnol. 40, 1023–1025. 10.1038/s41587-021-01156-3

Tooke, C.L., Hinchliffe, P., Bragginton, E.C., Colenso, C.K., Hirvonen, V.H.A., Takebayashi, Y., Spencer, J., 2019. β-Lactamases and β-Lactamase Inhibitors in the 21st Century. J. Mol. Biol., The molecular basis of antibiotic action and resistance 431, 3472–3500. 10.1016/j.jmb.2019.04.002

Uhde, K.B., Pathak, S., McCullum, I., Jannat-Khah, D.P., Shadomy, S.V., Dykewicz, C.A., Clark, T.A., Smith, T.L., Brown, J.M., 2010. Antimicrobial-resistant nocardia isolates, United States, 1995-2004. Clin. Infect. Dis. Off. Publ. Infect. Dis. Soc. Am. 51, 1445–1448. 10.1086/657399

Valdezate, S., Garrido, N., Carrasco, G., Villalón, P., Medina-Pascual, M.J., Saéz-Nieto, J.A., 2015. Resistance gene pool to co-trimoxazole in non-susceptible Nocardia strains. Front. Microbiol. 6, 376. 10.3389/fmicb.2015.00376

Wang, F., Cassidy, C., Sacchettini, J.C., 2006. Crystal Structure and Activity Studies of the Mycobacterium tuberculosis β-Lactamase Reveal Its Critical Role in Resistance to β-Lactam Antibiotics. Antimicrob. Agents Chemother. 50, 2762–2771. 10.1128/aac.00320-06

Wilson, J.W., 2012. Nocardiosis: updates and clinical overview. Mayo Clin. Proc. 87, 403–407. 10.1016/j.mayocp.2011.11.016

Zhao, P., Zhang, X., Du, P., Li, G., Li, L., Li, Z., 2017. Susceptibility profiles of Nocardia spp. to antimicrobial and antituberculotic agents detected by a microplate Alamar Blue assay. Sci. Rep. 7, 43660. 10.1038/srep43660

